# Alterations in the human plasma lipidome in response to Tularemia vaccination

**DOI:** 10.1101/2020.03.16.994525

**Authors:** Kristal M. Maner-Smith, David A. Ford, Johannes B. Goll, Travis L. Jensen, Manoj Khadka, Jennifer K Colucci, Casey E. Gelber, Carolyn J. Albert, Steve Bosinger, Jacob D. Franke, Muktha Natrajan, Nadine Rouphael, Robert Johnson, Patrick Sanz, Evan J. Anderson, Daniel F. Hoft, Mark Mulligan, Eric A. Ortlund

## Abstract

Tularemia is a rare but highly contagious and potentially fatal disease caused by bacteria *Francisella tularensis* where as few as ten inhaled organisms can lead to an infection, making it one of the most infectious microorganisms known and a potential bioweapon. To better understand the response to a live, attenuated tularemia vaccine and the biological pathways altered post-vaccination, healthy adults were vaccinated by scarification and plasma was collected pre- and post-vaccination for longitudinal lipidomics studies. Using tandem mass spectrometry, we identified and quantified individual lipid molecular species within representative lipid classes in plasma to characterize alterations in the plasma lipidome during the vaccine response. Separately, we targeted oxylipins, a subset of lipid mediators involved in inflammatory pathways. We identified 14 differentially abundant lipid species from eight lipid classes. These included 5-Hydroxyeicosatetraenoic acid (5-HETE), an eicosanoid produced following arachidonic acid liberation and epoxygenation, which is indicative of lipoxygenase activity and, subsequently, inflammation. Results suggest that 5-HETE was metabolized to a dihydroxyeicosatrienoic acid (DHET) by Day 7 post-vaccination, shedding light on the kinetics of the 5-HETE-mediated inflammatory response. In addition to 5-HETE and DHET, we observed pronounced changes in 34:1 phosphatidylinositol, anandamide, oleamide, ceramides, 16:1 cholesteryl ester, and several glycerophospholipids, several of these changes in abundance were correlated with serum cytokines and T cell activation. These data provide new insights into alterations in plasma lipidome post tularemia vaccination, potentially identifying key mediators and pathways involved in vaccine response and efficacy.

## Introduction

Vaccines initially trigger innate immune responses that result in development of an adaptive immune response with establishment of immunological memory (1). This process is incompletely understood but is probably best detailed for the Yellow Fever vaccine (2). This vaccine is highly effective and elicits a wide range of immune responses, resulting in greater than 30 years of protection from yellow fever. Previously published research has shown that the vaccine produces a biologic signature identified by transcriptomics, cytokine analyses and flow cytometry (3). Such biologic signatures may correlate with immunogenicity to predict the efficacy of novel vaccines. Lipids are key stress-response and immune signaling molecules and changes in their circulating concentrations reflect a host of immune processes(4) (5) (6). Thus, lipids play a critical role in inflammation. Accordingly, performing lipidomics analysis is a key element in the systems-wide multi-omic platform approaches leveraging data from transcriptomics, metabolomics, and proteomics to obtain a comprehensive picture of the vaccine response.

Tularemia, caused by the intracellular gram-negative bacterium *Francisella tularensis*, is transmitted from infected ticks and mosquitos to humans. *F. tularensis* has a mortality rate approaching 30% if untreated and is one of the most highly infectious bacteria known; infection with as few as 10 organisms can cause severe disease (7) (8, 9). Due to this high infectivity, *F. tularensis* could be weaponized and used as a biological weapon. A live attenuated tularemia vaccine has been used by the US Army for decades and can protect against severe disease. However, this vaccine is associated with significant reactivity and has not been widely used outside of at-risk populations. In addition, the US supply of this live-attenuated vaccine has been depleted. A prospective randomized study between a recently generated new and old lot of Tularemia vaccine showed similar immunogenicity (7). To evaluate the full complement of biological responses to this new lot of vaccine, we performed a multi-omics approach to discover changes in genome, proteome, metabolome, and lipidome(10) (11) (12). In this report, we explore changes in the plasma lipidome of vaccine recipients at Day 1, 2, 7, 14 post-vaccination as compared to lipidome pre-vaccination, with aims to determine lipid biomarkers of serological responses.

## Methods

### Study Design

For this study, a subset of plasma samples from a tularemia vaccine clinical trial were used (7). In original trial, 10 healthy subjects aged 18 to 45 years old were recruited and vaccinated with a single, undiluted dose of the *Francisella tularensis* live vaccine strain (LVS) lot 20 produced by DynPort Vaccine Company (DVC-LVS) via scarification on day 0. The vaccines were administered in the ulnar aspect of the volar surface (palm side) of the forearm midway between the wrist and the elbow (7). Here, we investigated plasma samples from 10 subjects taken pre-vaccination (Day 0) and at Days 1, 2, 7, 14 following Tularemia vaccination to assess response.

### Standards

Internal synthetic standards were obtained from Avanti Polar Lipids, Alabaster, AL. These include di-20:0 phosphatidylcholine (PC) (x:y where x indicates number of carbons and y indicates number of double bonds in fatty acid constituents), di-14:0 phosphatidylethanolamine (PE), N-17:0 sphingomyelin (SM), di-14:0 phosphatidylserine (PS), 17:0 lysophosphatidylcholine (LPC), 14:0 lysophosphatidylethanolamine (LPE), di-20:0 diacylglycerol (DAG), tri-17:0 triacylglycerol (TAG), and 17:0 free fatty acid (FFA), 17:0 cholesteryl ester (CE) and N-17:0 ceramide(Cer). Each of these internal standards were added at concentrations that approximate plasma concentrations in humans. Concentrations are reported in **Supplementary Table 1**.

### Lipid extraction

Targeted lipidomics experiments were conducted at two different mass spectrometry facilities using aliquots from the same subject and timepoint as a pilot of a larger consortia study. To reduce potential systematic errors, both laboratories performed the same lipid extraction technique and shared internal standards.

Plasma was prepared from whole blood samples using centrifugation and 100 ◻l of this plasma was extracted using a modified Bligh and Dyer lipid extraction (13). For this, 100 ◻L plasma was homogenized in 500 ◻l 2:1 v/v methanol:chloroform and vortexed to ensure homogeneity of sample. To this, 0.1mM sodium chloride was added to aide in the partitioning of zwitterionic lipids to the organic phase. The organic phase was then recovered, dried under nitrogen gas, and the lipid weight recorded. Recovered lipids were then reconstituted in 1 mL of 1:1 v/v chloroform:methanol prior to injection into the mass spectrometer.

To optimize PE ionization efficiency, an aliquot of extracted lipid was used to convert PE to PE-fMOC derivatives (14).

For oxylipin analysis, 200 ◻l plasma was extracted using C18 solid phase extraction (SPE) cassettes (15–18). Briefly, C18 cassettes were conditioned using ethyl acetate, methanol, and water, in succession. The sample was then deposited on the matrix and were rinsed with 3 column volumes of water and hexane, respectively. Oxylipins were then eluted with 3 column volumes of Methyl Formate. Recovered lipids were then dried and the lipid weight recorded. The samples were then suspended in 500 ◻L 1:1 chloroform:methanol.

### Mass Spectrometry

Targeted lipidomics was conducted using triple quadrupole mass spectrometers, SCIEX QTRAP5500 (Framingham, MA, USA) and a Thermo Quantum (Waltham, MA, USA) at EMO and SLU sites, respectively. Lipids were directly infused into the mass spectrometer. Instrumental parameters were optimized using the single, shared internal standard and were held constant during the course of the experiment. A table of instrumental parameters are shown in **Supplementary Table 2**.

To determine the distribution of lipids in extracted plasma samples, shotgun methodology was conducted whereby user specified lipid classes were selectively targeted using characteristic scans. PC species were quantified in the negative ion mode by neutral loss scanning at 50 amu, corresponding to the loss of methylene chloride from the chlorinated choline headgroup (19–23). Similarly, SM molecular species were quantified as chlorinated adducts in negative ion mode using neutral loss scanning of 50 amu (24). PE molecular species were quantified as their fMOC derivatives in negative ion mode using neutral loss scanning of 222.2 amu (14). Cholesteryl ester (CE) molecular species were quantified as sodiated adducts in the positive ion mode using neutral loss scanning of 368.5 amu as previously described (25). Ceramide (Cer) molecular species were quantified in negative ion mode using neutral loss scanning of 256.2 amu (26). A table of the targeted lipid classes and the scans used isolate these lipids is shown in **Table 1.** Characteristic transitions that are representative of key oxidized lipids were monitored and are presented in **Table 2**(27, 28).

**Table 1.** Table of characteristic scans. The following precursor ion scans were used to selectively target the associated lipid classes present in plasma samples.

| Lipid Class | Class Abbrev. | Ionization Mode | Characteristic Scan | Lipid IDs | Collision Energy |
| --- | --- | --- | --- | --- | --- |
| Phosphatidylethanolamine (Fmoc Deriv) | PE | Negative | NL222 | 35 | -30 |
| Phosphatidylcholine | PC | Negative | NL50 | 42 | -30 |
| Sphingomyelin | SM | Negative | NL50 | 4 | -24 |
| Phosphatidylinositol | PI | Negative | Prec241 | 9 | -30 |
| Phosphatidylserine | PS | Negative | NL87 | 1 | -30 |
| Cholesteryl Esters | CE | Positive | NL368 | 14 | 25 |
| Ceramide | Cer | Negative | NL256 | 11 | -32 |
| Oxidized Lipids | OXY | Negative | MRM† | 14 | -25 |

†: refer to Table 2

**Table 2.** Oxidized Lipids Panel Transition Table. A multiple reaction monitoring (MRM) based method was used to quantify the abundance of 14 oxidized lipid species. Species with the same transition and are grouped and reported together.

| Lipid ID | Lipid Species | Abbrev. | m/z | Transition |
| --- | --- | --- | --- | --- |
| TULIPID001 | Oleoylethanolamide | OEA | 325.5 | 325<62 |
| TULIPID002 | Arachidonoylethanolamine | AEA | 347.5 | 347<62 |
| TULIPID003 | Prostaglandin E2 Ethanolamide | PGE2 Ethanolamide | 395.5 | 378<62 |
| TULIPID004 | Prostaglandin F2a Ethanolamide | PGF2a Ethanolamide | 397.5 | 380<62 |
| TULIPID005 | **12(13)-EpOME/13-HODE** |  |  |  |
|  | *12(13)epoxy-9-octadecenoic acid* | *12(13)-EpOME* | 296.5 | 296<168 |
|  | *13-hydroxy-9,11-octadecadienoic acid* | *13-HODE* | 296.5 | 296<168 |
| TULIPID006 | 9,10-dihydroxy-12-octadecenoic acid | 9,10-DiHOME | 314.5 | 314<172 |
| TULIPID008 | Thromboxane B2 | TXB2 | 370.5 | 370<147 |
| TULIPID009 | 12-hydroxy-5,8,10,14-eicosatetraenoic acid | 12-HETE | 320.5 | 320<87 |
| TULIPID010 | **9-HETE/11(12)-EET** |  |  |  |
|  | *9-hydroxy-5,7,11,14-eicosatetraenoic acid* | *9-HETE* | 320.5 | 320<167 |
|  | *11(12)-epoxy-5,8,14-eicosatrienoic acid* | *11(12)-EET* | 320.5 | 320<167 |
| TULIPID011 | 20-hydroxy-5,8,11,14-eicosatetraenoic acid | 20-HETE | 320.5 | 320<289 |
| TULIPID012 | 5-hydroxy-6,8,11,14-eicosatetraenoic acid | 5-HETE | 320.5 | 320<301 |
| TULIPID013 | 8(9)-epoxy-5,11,14-eicosatrienoic acid | 8(9)-EET | 320.5 | 320<69 |
| TULIPID014 | 14(15)-epoxy-5,8,11-eicosatrienoic acid | 14(15)-EET | 320.5 | 320<220 |
| TULIPID007 | **Summed DHET Species** |  |  |  |
|  | *14,15-dihydroxy-5,8,11-eicosatrienoic acid* | *14,15-DHET* | 338.5 | 338<256,170,184,82 |
|  | *11,12-dihydroxy-5,8,14-eicosatrienoic acid* | *11,12-DHET* | 338.5 | 338<256,170,184,82 |
|  | *8,9-dihydroxy-5,11,14-eicosatrienoic acid* | *8,9-DHET* | 338.5 | 338<256,170,184,82 |
|  | *5,6-dihydroxy-8,11,14-eicosatrienoic acid* | *5,6-DHET* | 338.5 | 338<256,170,184,82 |

### Lipid quantification

Targeted lipids were quantified in fmol/*μ*L by dividing a sample’s peak intensity for a certain lipid by the corresponding internal control/standard lipid peak intensity and by multiplying it with the corresponding internal control/standard of known concentration. Corrections were also made for type I and type II ^13^C isotope effects (29). Additional corrections were made from response curves for CE molecular species (30). For oxylipins, peak intensity ratios were multiplied by the corresponding external control/standard’s molecular weight to obtain fmol/*μ*L. Standard lipidomics nomenclature was used throughout whereby acyl linkages are standard and ether linked lipids were denoted as : p=plasmalogen subclass and e=alkyl ether subclass. Aliphatic groups in lipid classes were also denoted as x:y where x= number of carbons and y= number of double bonds (e.g., 20:4 (arachidonic acid) has 20 carbons and 4 double bonds).

## Data analysis

### Missing Values and Baseline Calculations

Observations with 0 fmol/◻l were set to NA/missing and these values were imputed using the k-nearest neighbor algorithm implemented in the impute R package (Version 1.44.0). Lipids with at least 40/50 (80%) non-missing observations were used as input for imputation and downstream analysis. The number of neighbors to be used as part of the imputation step was set to 8. Subject-specific log2 lipid fold changes from baseline were calculated for each subject and post-vaccination day (day 1, 2, 7, 14) by subtracting baseline (day 0) imputed log_2_ fmol/◻L from each of the subject’s post-vaccination day imputed log_2_ fmol/◻L. After consolidating the data across experiments, the dataset included 129 unique lipids of which 116 had at least 80% non-missing observations. The complete list of identified lipids is provided in **Supplementary Table 3**.

### Statistics

Data was analyzed using the R statistical programming language (Version 3.2.5) and R Bioconductor.

Lipids that significantly differed in their response from baseline were identified by using a two-sided permutation paired t-test comparing baseline (day 0) to post-vaccination (day x) lipid fmol/ (H0 : u_(dayx - day0)_ = 0, H1 : u_(dayx - day0)_ ≠ 0; on the log_2_ scale). We used a combination of permutation p-value cut off of < 0.05 and an effect size cut off of ≥ 1.2-fold difference between mean pre- and post-vaccination responses to determine significantly differentially abundant (DA) lipids for each post-vaccination day.

Confidence intervals of the median fold change response for the time trend plots were calculated using bootstrapping (1,000 samples each).

### Correlation Networks

Associations between DA lipid log_2_ fold changes and cytokine log_2_ fold changes or peak T-cell and tularemia-specific microagglutination responses were assessed using Spearman correlation and visualized using the R igraph package. Peak T-cell responses (CD3+CD4+CD38+HLA-DR+ cells and CD3+CD8+CD38+HLA-DR+ cells) were based on the peak percentage of activated cells post-vaccination at Days 7, 14, or 28. Peak tularemia-specific microagglutination titer was represented as the log_2_ transformed peak titer observed at Days 14 or 28(7).

## Results

Extracted lipids were analyzed by tandem mass spectrometry, using both traditional shotgun lipidomics as well as MRM-based methods. A series of characteristic scans were performed to isolate each abundant lipid class present in plasma (**Table 1–2**). Overall, 129 lipids were identified (Supplemental Table S2), of which 116 lipids, consisting of 14 oxylipins, 75 phospholipids, 14 cholesteryl esters, and 13 sphingolipids, had sufficient non-missing data to be analyzed. Fourteen of the 116 lipids were found to be DA (p < 0.05, fold change ≥ 1.2) at Day 1, 2, 7, or 14 post vaccination compared to pre-vaccination (Day 0) and are presented in **Table 3**. A breakdown of these lipids according to classification is shown in **Figure 1.** Phospholipids showed the largest proportion of DA (50.0%), followed by oxylipins (28.6%), cholesteryl esters (14.3%), and sphingolipids (7.7%).

**Figure 1.**
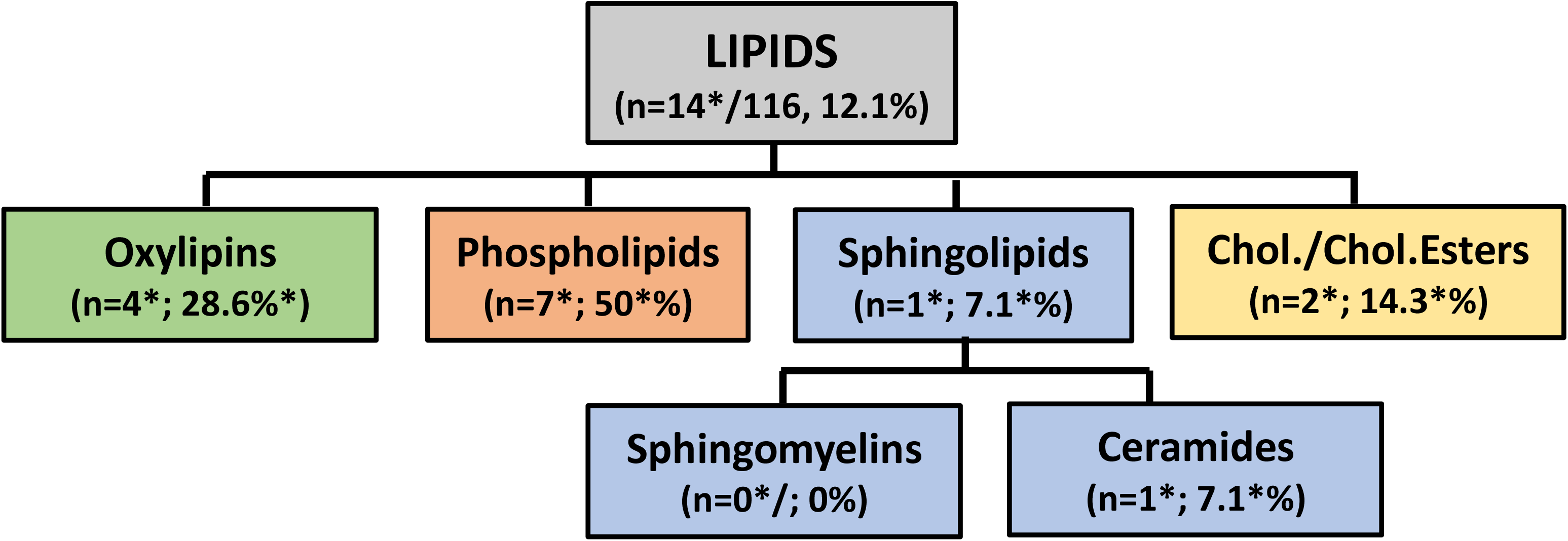
Schematic of lipid classes identified by targeted lipidomics. The following lipid classes and associated number of lipids were quantified in patient plasma samples. The asterisk indicates the number of lipids that showed a differential response relative to pre-vaccination following vaccination. Phospholipids contained the largest proportion of differentially abundant lipids (7 DA lipids, 50%), followed by oxidized lipids (4 DA lipids, 28.6%), cholesterol esters (2 DA lipids, 14.3%), and sphingolipids (1 DA lipid, 7.1%).

**Table 3.** Overview of differentially abundant lipids by day. A two-sided permutation paired t-test was used to determine lipids with post-vaccination plasma molar concentrations that significantly differed compared to pre-vaccination. Lipids with a p-value < 0.05 and an increase/decrease from pre-vaccination of at least 20% (◻1.2-fold) were deemed differentially abundant.

| Lipid Name | Lipid ID | Lipid Class | Day | Fold Change | t-statistic | p-value |
| --- | --- | --- | --- | --- | --- | --- |
| 16:1 LPC | TULIPID017 | PC | Day 1 | 0.709 | -2.7 | 0.0137 |
| 20:3 LPC | TULIPID024 | PC | Day 1 | 0.74 | -5.5 | 0.002 |
| 36:3 PE | TULIPID103 | PE | Day 1 | 1.238 | 1.8 | 0.0469 |
| p40:6 PE | TULIPID119 | PE | Day 1 | 1.214 | 2.4 | 0.0449 |
| 16:1 CE | TULIPID078 | CE | Day 2 | 0.811 | -4.7 | 0.0039 |
| 16:1 LPC | TULIPID017 | PC | Day 2 | 0.597 | -2.9 | 0.0234 |
| 20:3 LPC | TULIPID024 | PC | Day 2 | 0.62 | -3.8 | 0.0078 |
| 22:3 CE | TULIPID089 | CE | Day 2 | 0.376 | -2.6 | 0.0371 |
| 34:1 PI | TULIPID058 | PI | Day 2 | 0.388 | -2.7 | 0.0254 |
| 34:2 PE | TULIPID099 | PE | Day 2 | 0.797 | -3.4 | 0.0098 |
| AEA | TULIPID002 | OXY | Day 2 | 0.686 | -2.7 | 0.0312 |
| OEA | TULIPID001 | OXY | Day 2 | 0.682 | -3.0 | 0.0156 |
| 16:1 CE | TULIPID078 | CE | Day 7 | 0.824 | -2.2 | 0.0371 |
| 16:1 LPC | TULIPID017 | PC | Day 7 | 0.534 | -2.6 | 0.0312 |
| 20:3 LPC | TULIPID024 | PC | Day 7 | 0.598 | -3.0 | 0.0195 |
| 22:0 C | TULIPID072 | CE | Day 7 | 0.807 | -2.9 | 0.0254 |
| 38:3 PI | TULIPID064 | PI | Day 7 | 0.491 | -2.4 | 0.0469 |
| 5-HETE | TULIPID012 | OXY | Day 7 | 0.386 | -2.7 | 0.0312 |
| Summed DHET Species | TULIPID007 | OXY | Day 7 | 1.969 | 3.1 | 0.0215 |
| 5-HETE | TULIPID012 | OXY | Day 14 | 0.674 | -2.3 | 0.0488 |

To investigate changes over time for DA lipids, we visualized DA lipid responses from pre-vaccination using radar plots. **Figure 2** is a radar plot showing the magnitude of change of the 19 differentially abundant lipids identified. Summed DHET species showed the largest increase in molar concentration compared to pre-vaccination overall with a 2-fold increase over pre-vaccination at Day 7 post-vaccination. By contrast, 5-HETE showed diametrically opposed changes at Day 14 following tularemia vaccination.

**Figure 2.**
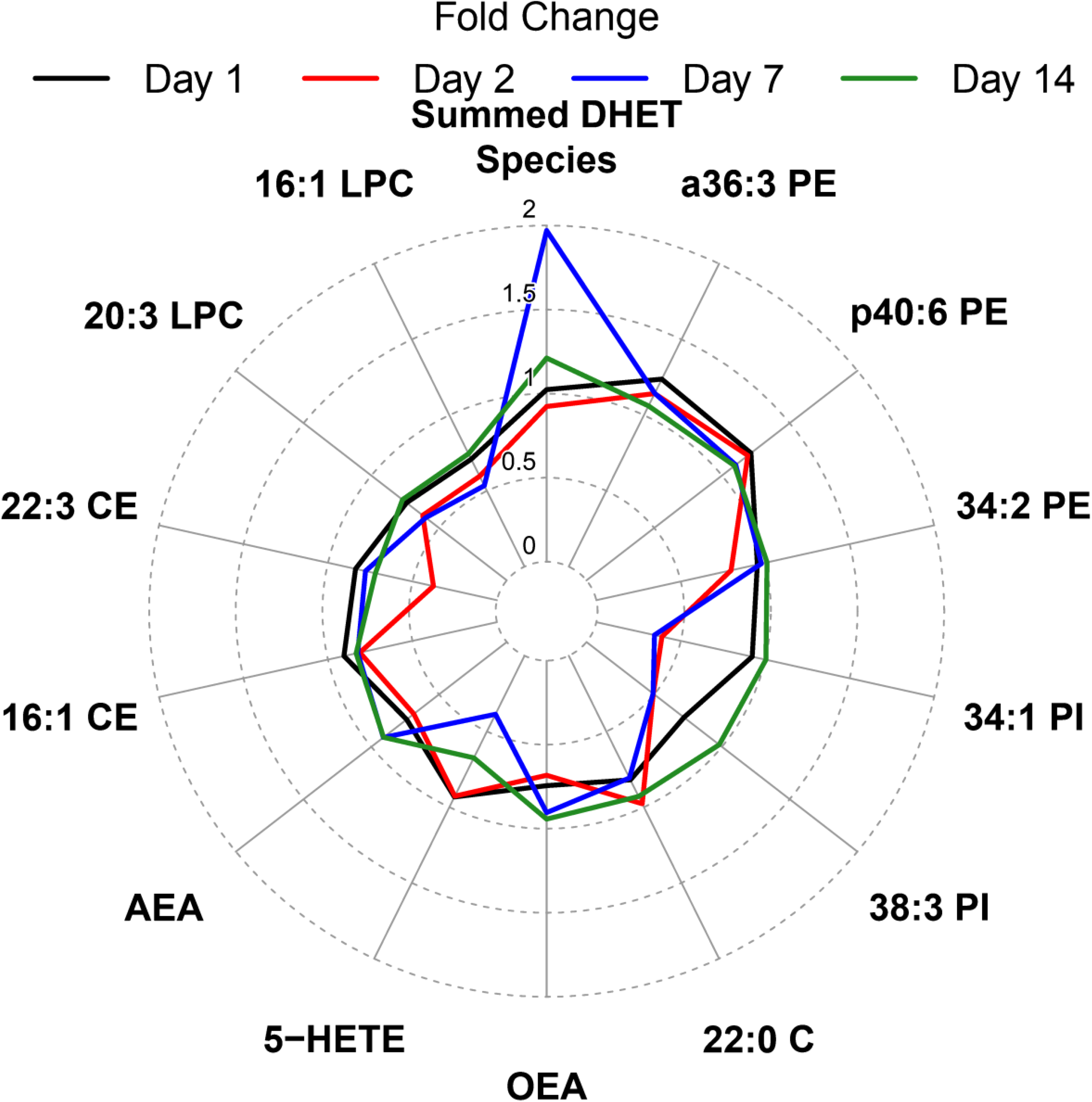
Radar plot summarizing mean fold change from pre-vaccination by day and DA lipids. Lines represent the magnitude of the mean fold change of differentially abundant lipid species at Day 1, 2, 7, and 14 compared to pre-vaccination. Lipids are ordered clockwise by descending maximum absolute log_2_ fold change starting at the top center.

To further investigate changes in DHET species in relation to 5-HETE, we plotted log_2_ fold change time trends and associated 95% confidence intervals over time (Figure 3). **Panel 3A** outlines the mechanism whereby 5-HETE is converted to DHET through the enzymatic action of the enzyme encoded by the *CYP4F3* gene, while **Panels 3B and 3C** show inverse fold changes of both species showing a maximum decrease for 5-HETE and a maximum increase for DHET at Day 7 post vaccination and returning to close to baseline levels at Day 14.

**Figure 3.**
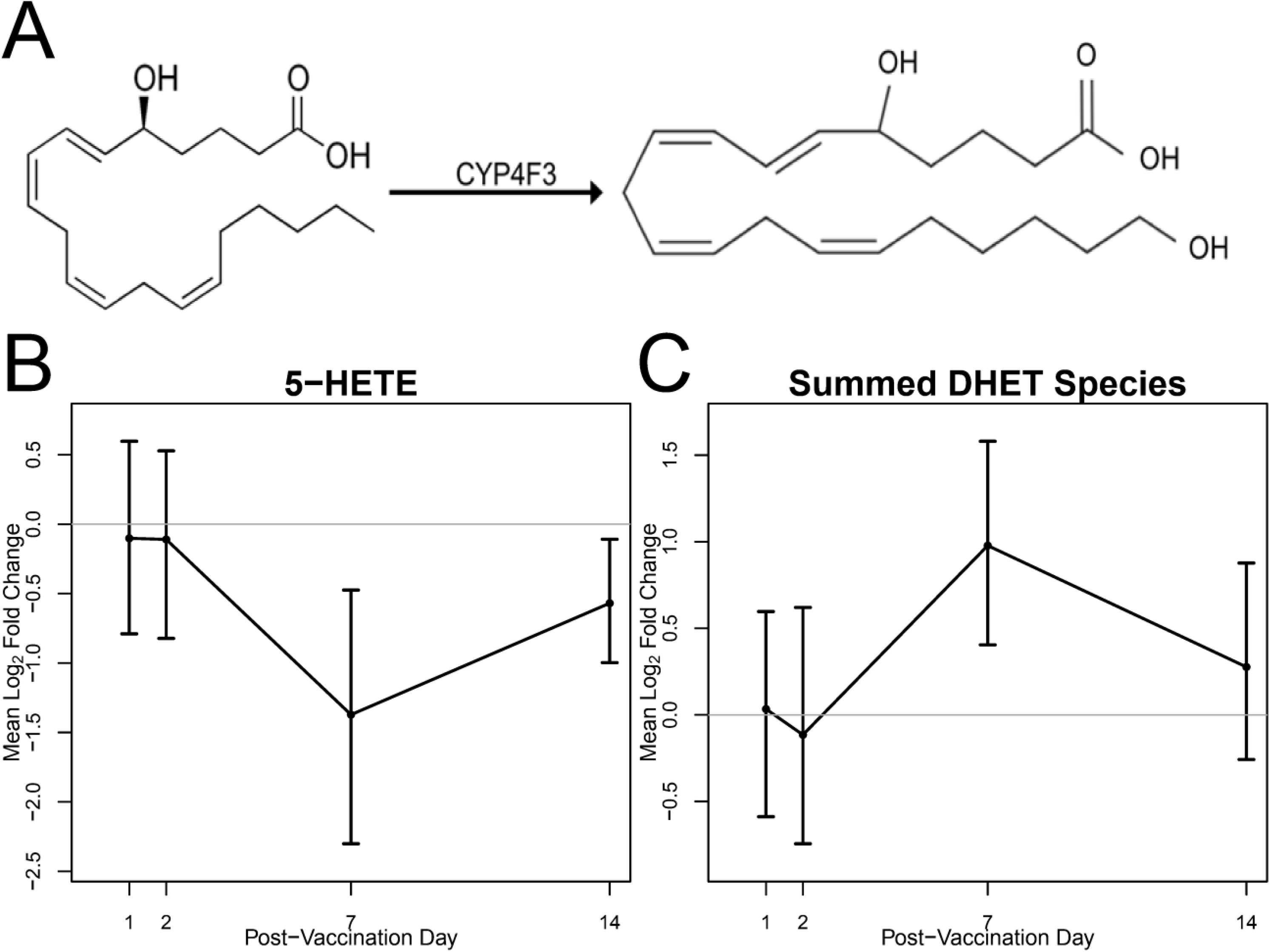
5-HETE is metabolized by CYP4F3 to form 5,20-DHET. A) 5-HETE is converted to DHET species by CYP4F3. B and C) 5-HETE and DHET Time trends of mean fold change from pre-vaccination and associated 95% bootstrap CIs. The abundance of 5-HETE and combined DHET species is found to be differentially abundant at 7 and 14 days post vaccination.

In addition to bioactive oxylipins, CE abundance was evaluated. **Figure 4** is a time-trend plot that probes the plasma abundance of DA lipid 16:1CE versus transcript of ABCA1, a protein target responsible for cholesterol export. The abundance of 16:1CE decreased at Day 2 post vaccination. In addition, the *ABCA1* gene encoding for the primary protein responsible for cholesterol efflux from macrophages and vascular tissue to circulating high density lipoprotein, showed a significant decrease at all post-vaccination days except Day 2 demonstrating that cholesterol homeostasis mechanisms may also be affected by tularemia vaccination.

**Figure 4.**
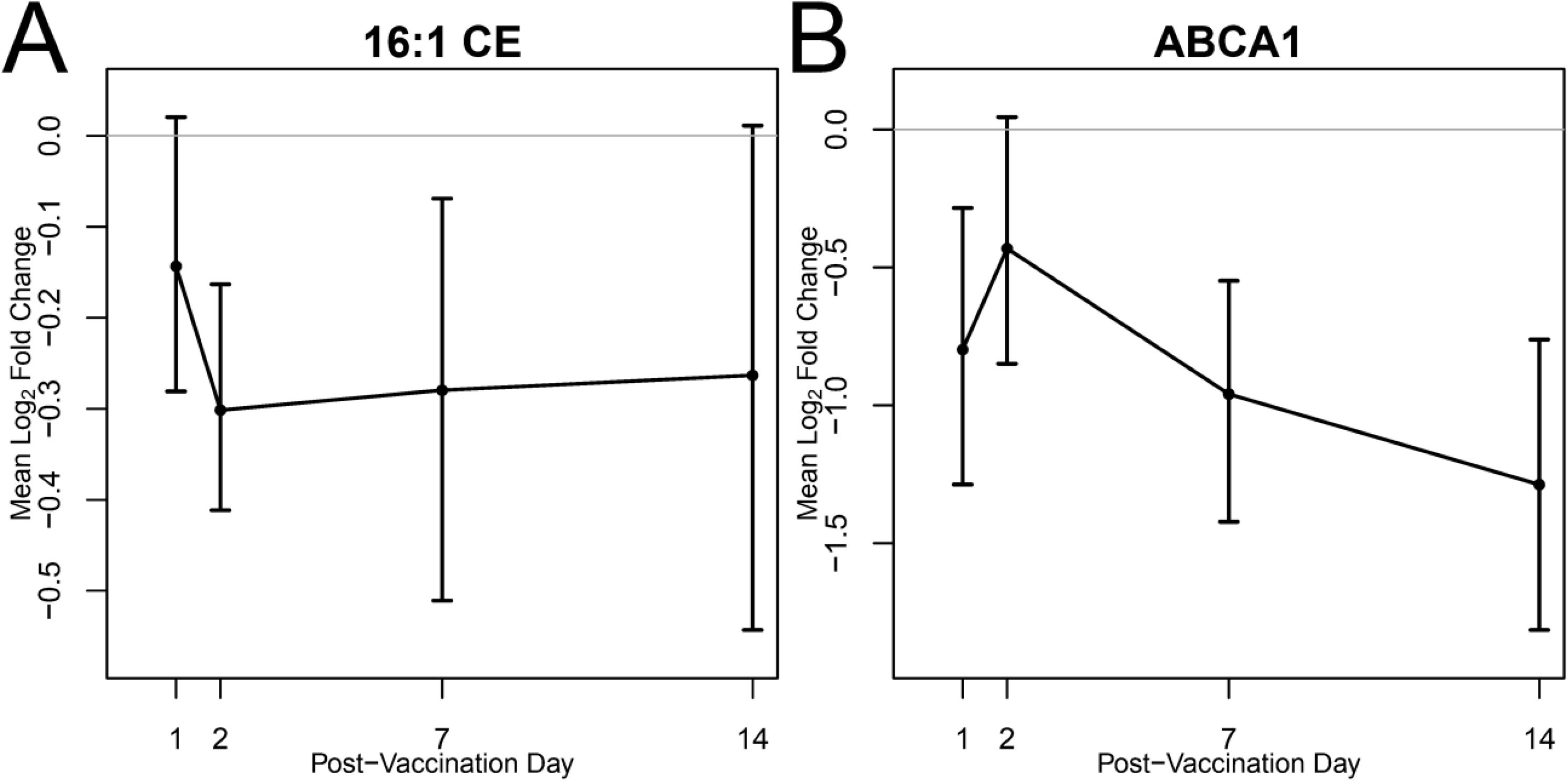
Reduction in cholesterol esters compared to pre-vaccination. A, B: Time trends of mean log_2_ fold change from pre-vaccination and associated 95% bootstrap CIs. Fold change of A) 16:1 CE molar concentration and B) ABCA1 gene expression relative to pre-vaccination. Gene expression of the ABCA1 gene was significantly decreased at all post-vaccination days but Day 2.

The 50% of the identified DA lipids were phospholipids, some containing esterified biologically active lipids such as arachidonic acid (20:4) and oleic acid (18:1) (**Figure 5**). These lipids are involved in inflammatory pathways leading to the conversion of 20:4 and 18:1 to endocannabinoids AEA and OEA, respectively, as outlined in Figure **5A**. The plasma abundance of these lipids (**Panel 5b**) reveals that both OEA and AEA decrease at Day 2 post vaccination and returning to baseline by Day 14.

**Figure 5.**
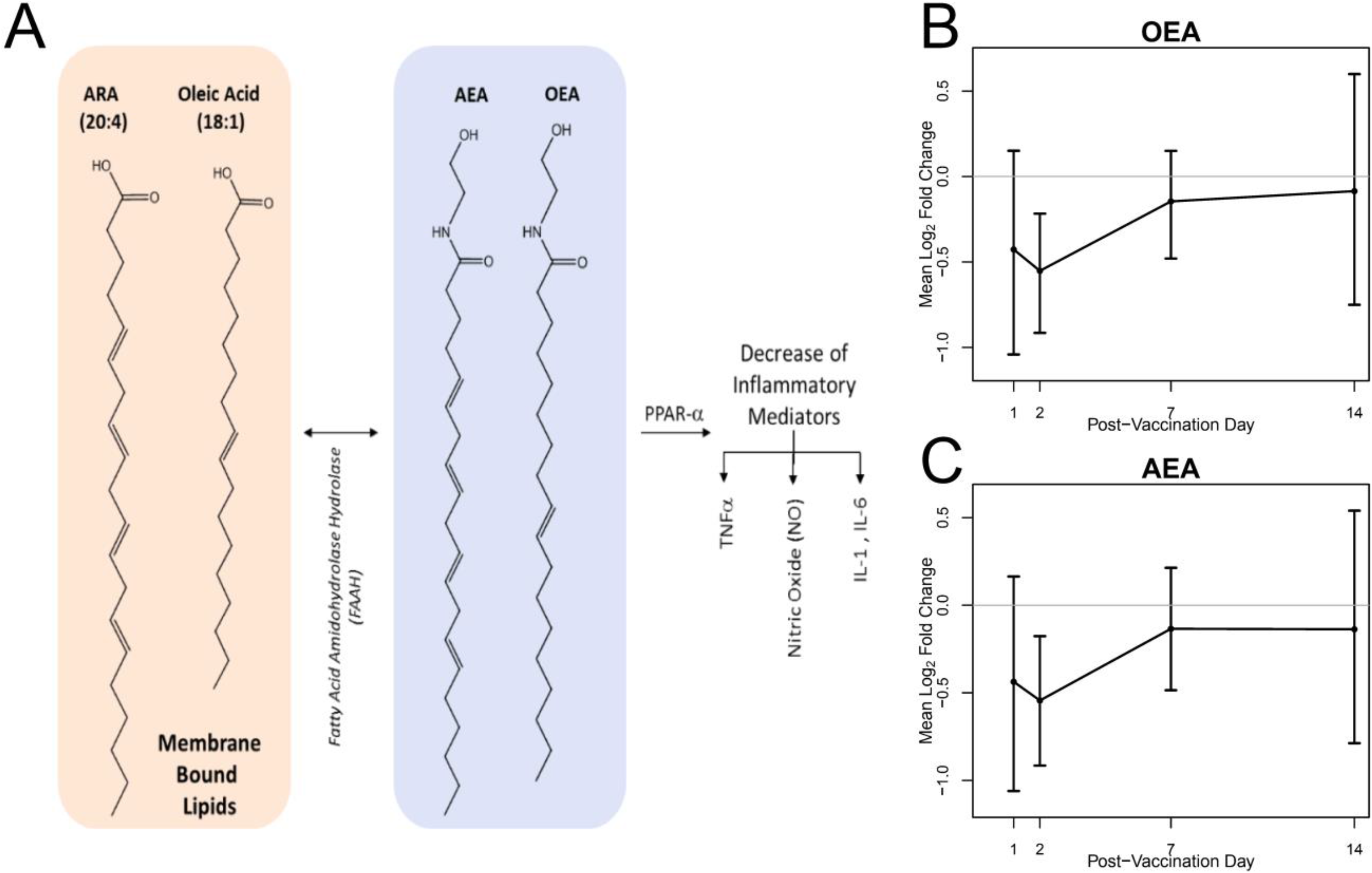
Biosynthesis of AEA and OEA from membrane lipids and plasma abundance. A) AEA and OEA are synthesized from membrane bound lipids and are activators of PPAR-a, whose activation decreases cellular abundance of inflammatory mediators. B and C) OEA and AEA Time trends of mean fold change from pre-vaccination and associated 95% bootstrap CIs. The plasma molar concentration of these species were found to be differentially abundant at Day 2 when compared to pre-vaccination.

To further characterize the role of DA lipids in the vaccine response, we generated a correlation network between log_2_ fold changes of 14 DA lipids, 22 cytokines, and CD4/CD8 T lymphocytes were assessed using Spearman correlation (**Figure 6**). Serum cytokines included EOTAXIN, basic fibroblast growth factor (FGF-BASIC), granulocyte colony-stimulating factor (G CSF), granulocyte-macrophage colony stimulating factor (GM CSF), interferon gamma (IFN-γ), interleukin 10 (IL-10), interleukin 12 p70 (IL-12 P70), interleukin 13 (IL-13), interleukin 17 (IL-17), interleukin 1 receptor agonist (IL-1RA), interleukin 2 (IL-2), interleukin 4 (IL-4), interleukin 5 (IL-5), interleukin 8 (IL-8), interleukin 9 (IL-9), interleukin 10 (IP-10), monocyte chemoattractant protein-1/monocyte chemotactic and activating factor (MCP-1/MCAF), macrophage inflammatory protein 1b (MIP 1B), platelet derived growth factor BB (PDGF BB), RANTES, tumor necrosis factor alpha (TNF-◻◻, and vascular endothelial growth factor (VEGF).

**Figure 6.**
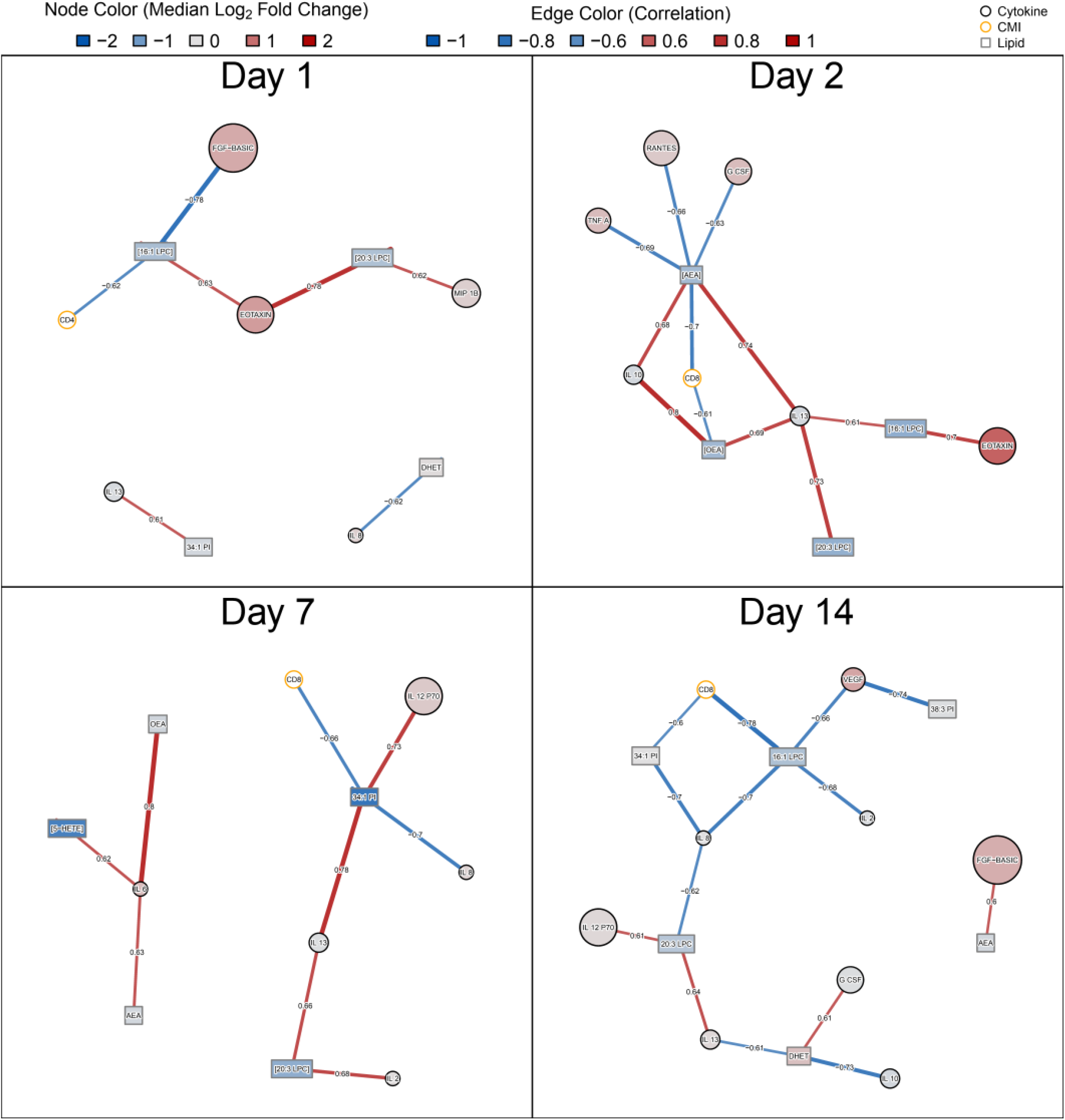
Correlation network summarizing associations between DA lipid fold changes and changes in serum cytokines as well as peak T-cell activation. Pairwise Spearman correlations were assessed between fold changes of 14 DA lipids and 22 serum cytokines by post-vaccination day. In addition, correlations with peak CD4+ and CD8+ T-cell activation at Days 7, 14, or 28 and tularemia-specific microagglutination were assessed. Black nodes represent lipids (rectangles) and cytokines (circles) while edges represent the Spearman correlation between fold changes. T-cell activation variables are shown in orange circles. Cytokine and lipid nodes are color-coded by log_2_ fold change. Edges are color coded and edge-widths are scaled by Spearman correlation. To facilitate visual interpretation, networks were filtered for Spearman correlation > 0.6.

On Day 1, CD4+ T-cell activation was negatively correlated with change in molar concentration of 16:1 LPC which was decreased (**Figure 6**). At Day 2, changes in molar concentration of OEA and AEA in plasma correlated with anti-inflammatory cytokines IL-10 and IL-13 and CD8 lymphocytes. In contrast, changes in OEA and AEA were negatively correlated with later peak CD8+ T-cell activation (**Figure 6)**. At Day 7, changes in OEA and AEA were associated with pro-inflammatory cytokine IL-6. The decrease in 5-HETE was positively associated with IL-6 as well. By Day 7, CD8 was negatively associated with 34:1 PI and any associations with OEA and AEA were no longer seen. At Day 14, change in DHET abundance was negatively associated with anti-inflammatory cytokines IL-10 and IL-13 and positively correlated with G CSF (**Figure 6**). 16:1 LPC showed a negative correlation with CD8+ cell activation. No correlation with CD4+ T-cell activation or microagglutination titer was observed with Spearman correlation ≥ 0.6 (7).

## Discussion

Lipids are a large and diverse family of molecules with considerable structural and biological diversity. The abundance of several integral classes of lipids is differentially affected by administration of the Tularemia vaccine. Loss of CE may indicate diminished hepatic cholesterol biosynthesis, which is one of the major contributors to plasma cholesterol levels carried by lipoproteins (31–33). TLR-signaling mediated changes in immune cell cholesterol metabolism (34) may explain some of the reduced CE signal as well: “on a cellular level, activation of TLR signaling leads to decreased cholesterol efflux, which results in further cholesterol accumulation and the amplification of inflammatory responses” (35). TLR signaling was enriched in DE gene responses at Day 2 following vaccination with increased expression for genes encoding for TLR5 (flagellin pattern recognition) and TLR6 (microbe-associated molecular pattern recognition) receptors proteins. The *ABCA1* gene encoding for the protein that exports excess cholesterol from cells was indeed significantly down-regulated in the transcriptomics data at Days 1, 7, and 14 with decreasing expression over time (43%, 48%, and 59% reduction, respectively, **Figure 4B**).

In addition CEs, the plasma concentrations of select oxylipins were also differentially abundant. We observed statistically significant changes in plasma levels of oxylipins as well as their associated precursors and inactive metabolites. The increased metabolism (decreased abundance) of the pro-inflammatory 5-HETE and the increase in DHET levels (inactive or less active metabolites of 5-HETE) suggest a role of these lipids by day 7, suggestion resolution of the inflammatory response mediated by this class of molecules (**Figure 3**). By day 7 (with conversion to DHET), the abundance of 5 HETE had fallen to 61% of the level present pre-vaccination, suggesting a negative regulation. We hypothesize this followed an increased abundance of this pro-inflammatory molecule as part of the innate response. These data are corroborated by transcriptomics data, which demonstrated that gene expression for gene *CYP4F22* was upregulated on day 2 post-vaccination (1.7-fold increase), day 7 post-vaccination (1.6-fold increase), and day 14 post-vaccination (1.8-fold increase) compared to pre-vaccination (data not shown). Together, these data suggest a clearing of acute inflammatory molecules via break down to their less active metabolites and thus a termination of the innate immune response. 5-Hydroxyeicosatetraenoic acid (5-HETE) is an eicosanoid that acts as an autocrine and paracrine signaling agent that contributes to the up-regulation of acute inflammatory and allergic responses. 5-HETE and its stereoisomers stimulate cells through the binding and activation of the GPCR oxoeicosanoid receptor 1 (OXER1). In turn, OXER1 activates the MAPK/ERK pathway, p38 mitogen-activated protein kinases, cytosolic phospholipase A2, PI3K/AKT, protein kinase C ß/ Ɛ, and ionic calcium channels. 5-HETE stimulates its target cells to degranulate in order to release anti-bacterial cytokines, produce bactericidal and tissue-injuring ROS, and activate the functions of the innate immune system. 5-HETE has also been shown to activate Peroxisome Proliferator-Activated Receptor (PPAR) isoform gamma, which is expressed in macrophages and dendritic cells and plays an important role in resolving inflammation (36) (37). Transcriptomics responses as measured using RNA-Seq indicated increasing pathway enrichment results for several published PPAR◻ -related Immunologic Signature Gene Sets including GSE25123 (gene expression signals in macrophage-specific PPAR◻ knockout mice) (38) with increasing enrichment signals over time (15, 17, 39, and 51 overlapping DE genes for Days 1, 2, 7, and 14, respectively).

The lifetime of 5-HETE is regulated through enzymatic conversion to inactive or less active metabolites, dihydroxyeicosatetraenoic acids (DHETs or diHETEs) by the Cytochrome P450F family of proteins including *CYP4F3* (**Figure 3**). DHET’s are derived from arachidonic acid (ARA) and are created from the hydrolysis, epoxygenation and, subsequent reduction of ARA from membrane lipids as a part of the cytochrome P450 epoxygenase pathway. The upregulation of this lipid has been shown in patient samples who have high degrees of meta-inflammation. DHETs have also been shown to contribute to coronary vasodilation in micromolar (non-physiological) concentrations.

In addition to 5-HETE, two additional differentially abundant oxylipins were identified over the time course of the study. These are OEA and AEA (**Figure 5**). Both are endogenous ligands of PPAR◻, a widely expressed transcription factor whose ligand-mediated activation inhibits increases in proinflammatory mediators such as TNF-a, IL-1b, IL-6 and others (39) (37). OEA, the most potent of these, is synthesized in enterocytes and begins with the N-acylation of oleic acid (18:1) from membrane PCs, such as those measured in this study, i.e. 34:1 PC and 36:1 PC (TULIPID038 and TULIPID044, respectively). These PC species were quantified but not determined to be differentially abundant (data not shown). It should be noted, however, that these lipids are major components of biomembranes and large changes in their abundance cannot be supported by the membrane, suggesting a feedback mechanism to support the synthesis of this lipid from dietary lipids.

Like OEA, AEA is also synthesized from membrane lipids. Membrane PE species that contain arachidonate at the sn-2 position and a saturated tail, such as palmitate or stearate, at the sn-1 position are acted upon by several enzymes in distinct pathways to liberate N-acylated arachidonate, AEA. AEA is a potent activator of the cannabinoid receptor as well as PPAR both receptors have well documented roles in inflammation [REFS]. In the present study, the plasma concentrations of both OEA and AEA significantly increase at days 7 and 14 in comparison to day 0. Combined, this suggests a resolution of inflammation and is consistent with changes in the abundance of 5-HETE.

Overall, this analysis showed that vaccination results in significant changes in the lipidome and that peak changes are observed seven days post vaccination. Lipids with the greatest changes play a role in mediating inflammation. Moreover, oxylipin data suggests a clearing of acute inflammatory molecules (5-HETE) via break down to their less active metabolites (DHET) and thus a termination of the innate immune response at Day 7. Similarly, decreased levels of cholesteryl esters accompanied by down-regulation of ABCA1 gene expression indicates pro-inflammatory pathways may be involved including mechanisms for cholesterol removal via the ABCA1 transporter. This was corroborated by the correlation with the abundance of key pro-inflammatory cytokines as shown in **Figure 6**. There was an inverse correlation between abundance of the pro-inflammatory TNF-◻ cytokine and AEA on Day 2. Similarly, on Day 2 there is a negative correlation between AEA and CD8+ T cells, implying heightening of immune response post vaccination (40) (40, 41). Similarly, an increase in the abundance of OEA occurs with a decrease in the pro-inflammatory IL-6 cytokine on Day 7, is in line with prior studies and suggesting the resolution of inflammation(42, 43) (39, 43). While no associations were identified between DA lipid responses and tularemia-specific microagglutination titer, CD4+ T-cell activation was linked to DA lipids at Day 1 while CD8 T-cell activation was associated with DA lipid changes at Days 2, 7, and 14. Since there is no association between lipids and microagglutination, this implies that lipids are involved in the cell specific signaling of T cells, supported by separate findings (44).

Taken together, we report novel results that cholesterol esters, oxylipins and certain PE species were differentially abundant post tularemia-vaccination, especially during the innate immune phases of vaccine response, suggesting that these lipids may be useful as future biomarkers of serological response to vaccination.

## Conflict of Interest Statement

EJA has received grant support through his institution from MedImmune, Pfizer, Merck, Sanofi Pasteur, PaxVax, Novavax, GSK, PaxVax, and Micron Biomedical, and consulting fees from AbbVie and Pfizer. All other authors do not have a commercial or other association that might pose a conflict of interest.

## Acknowledgements

We thank the Emory VTEU administrative and finance core for their support including Dean Kleinhenz, Hannah Huston, and Michele Paine McCullough.

## Funding Statement

This work was supported by awards from the Division of Microbiology and Infectious Diseases, National Institute of Allergy and Infectious Diseases at the National Institutes of Health to the Emory Vaccine and Treatment Evaluation Unit, contracts HHSN272200800005C and HHSN272201300018I, St. Louis University contract HHSN2722013000021I, and EMMES contract HHSN272201500002C. We also acknowledge the support of the Georgia Research Alliance to The Hope Clinic. This study was supported in part by the Emory Integrated Lipidomics Core (EILC), which is subsidized by the Emory University School of Medicine and is one of the Emory Integrated Core Facilities. Additional support was provided by the Georgia Clinical & Translational Science Alliance of the National Institutes of Health under Award Number UL1TR002378.

---

**Supplementary Table 1.** Table of Lipidomics Standards. The following synthetic standards were used to optimize instrumental parameters and also for quantification of patient plasma samples.

| OXY Standards |  |  | Shotgun Standards |  |  |
| --- | --- | --- | --- | --- | --- |
| Lipid Abbrev. | Lipid Species | Conc. (pg/ml) | Lipid | Lipid Class | [Conc; ug/ul] |
| OEA | Oleoylethanolamide | 14.2 |  |  |  |
| AEA | Arachidonoylethanolamine | 28.4 | di20:0 PC | Phosphatidylcholine | 60.0 |
| PGE2 EA | Prostaglandin E2 Ethanolamide | 11.4 | di14:0 PE | Phosphatidylethanolamine | 1.0 |
| PGF2a EA | Prostaglandin F2a Ethanolamide | 14.2 | 17:0 SM | Spingomyelin | 4.0 |
|  | 12(13)-EpOME/13-HODE |  |  |  |  |
| 12(13) EpOME | 12(13)epoxy-9-octadecenoic acid | 1.1 | di20:0 DG | Diacylglycerol | 4.0 |
| 13 HODE | 13-hydroxy-9,11-octadecadienoic acid | 1.4 | 17:0 FFA | Free Fatty Acid | 1.0 |
| 9,10 DIHOME | 9,10-dihydroxy-12-octadecenoic acid | 1.1 | Tri17:1 TG | Triacylglycerol | 10.0 |
| TBX2 | Thromboxane B2 | 11.4 | di14:0 PS | Phosphatidylserine | 0.1 |
| 12 HETE | 12-hydroxy-5,8,10,14-eicosatetraenoic acid | 1.1 | 17:0 LPC | Lysophosphatidylcholine | 2.5 |
|  | 9-HETE/11(12)-EET |  | 14:0 LPE | Lysophosphatidylethanolamine | 0.1 |
| 9 HETE | 9-hydroxy-5,7,11,14-eicosatetraenoic acid | 1.1 |  |  |  |
| 11(12) EET | 11(12)-epoxy-5,8,14-eicosatrienoic acid | 1.1 |  |  |  |
| 20 HETE | 20-hydroxy-5,8,11,14-eicosatetraenoic acid | 1.1 |  |  |  |
| 5 HETE | 5-hydroxy-6,8,11,14-eicosatetraenoic acid | 1.1 |  |  |  |
| 8(9) EET | 8(9)-epoxy-5,11,14-eicosatrienoic acid | 1.1 |  |  |  |
|  | 14(15)-epoxy-5,8,11-eicosatrienoic acid | 1.1 |  |  |  |
|  | Summed DHET Species |  |  |  |  |
| 14,15 DHET | 14,15-dihydroxy-5,8,11-eicosatrienoic acid | 1.1 |  |  |  |
| 11,12 DHET | 11,12-dihydroxy-5,8,14-eicosatrienoic acid | 1.1 |  |  |  |
| 8,9 DHET | 8,9-dihydroxy-5,11,14-eicosatrienoic acid | 1.1 |  |  |  |
| 5,6 DHET | 5,6-dihydroxy-8,11,14-eicosatrienoic acid | 1.1 |  |  |  |

**Supplementary Table 2.** Table of instrumental parameters. A list of optimized instrumental parameters used to conduct targeted lipidomics studies at EMO and SLU laboratories, respectively.

| EMO Instrumental Parameters |  |  |  | SLU Instrumental Parameters |  |  |  |
| --- | --- | --- | --- | --- | --- | --- | --- |
| Curtain Gas (CUR) | 25.00 | Collision Gas (CAD) | Low | Curtain Gas (CUR) | 20.00 | Sheath Gas | 12 arb. units |
| ESI Voltage (IS) | -3500 | Gas Source 1 (GS1) | 55.00 |  |  |  |  |
| ESI Temp (TEM) | 650°C | Gas Source 2 (GS2) | 50.00 | ESI Voltage (IS) | 3500 | Ion Sweep Gas | 1 arb. Units |
| Declustering Potential (DP) | -90.00 | Collision Energy (CE) | -40.00 |  |  |  |  |
| Entrance Potential (EP) | -10.00 | Collision Energy Spread (CES) | 0.00 | Capillary Temp. | 270°C | Auxillary Gas | 2 arb. units |
| Q1 and Q3 Resolution | Unit | Step Size | 0.2Da | Collision Pressure (Argon) | 1 atm | Capillary Offset | 35 |

**Supplementary Table 3.**
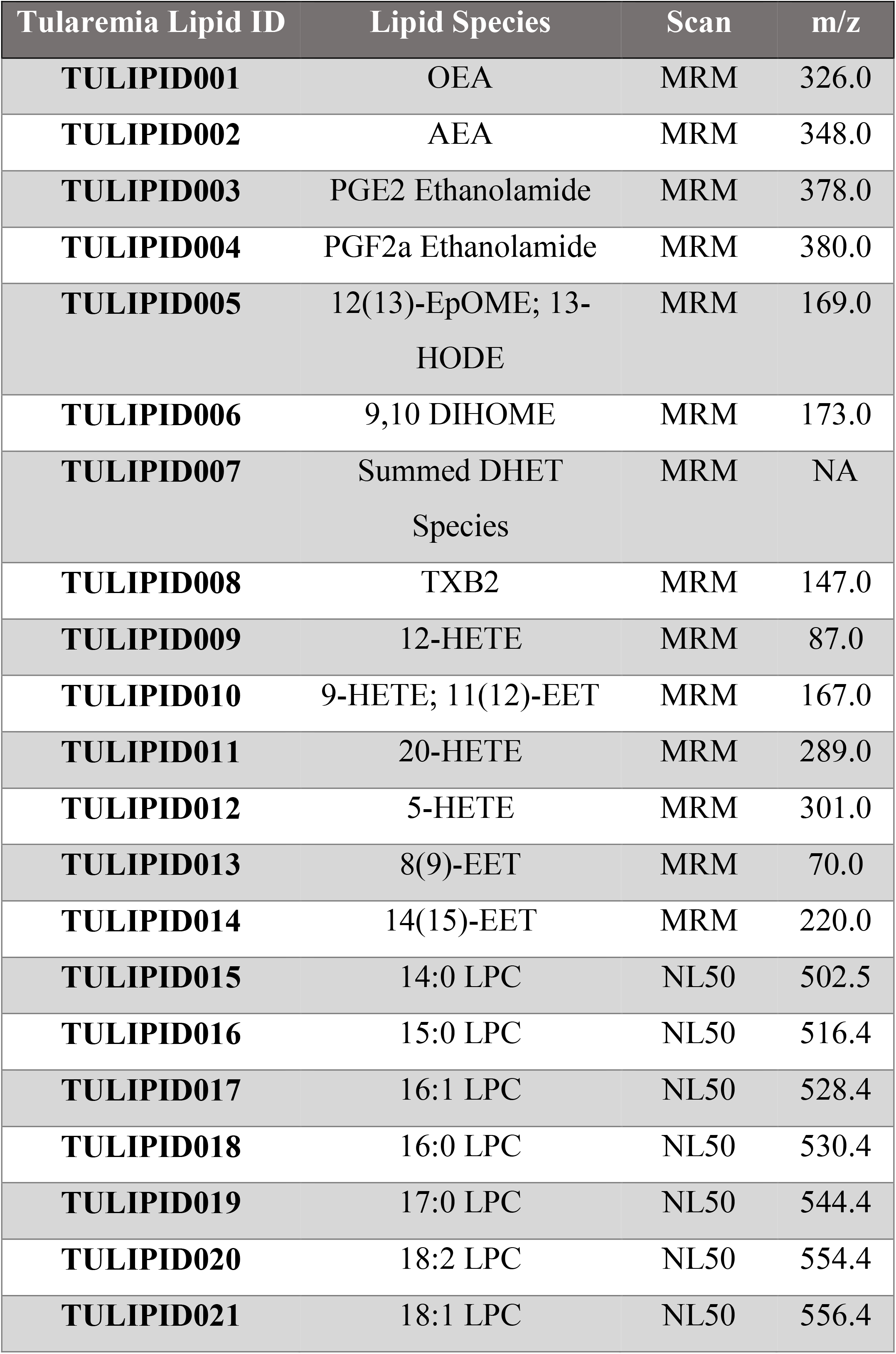
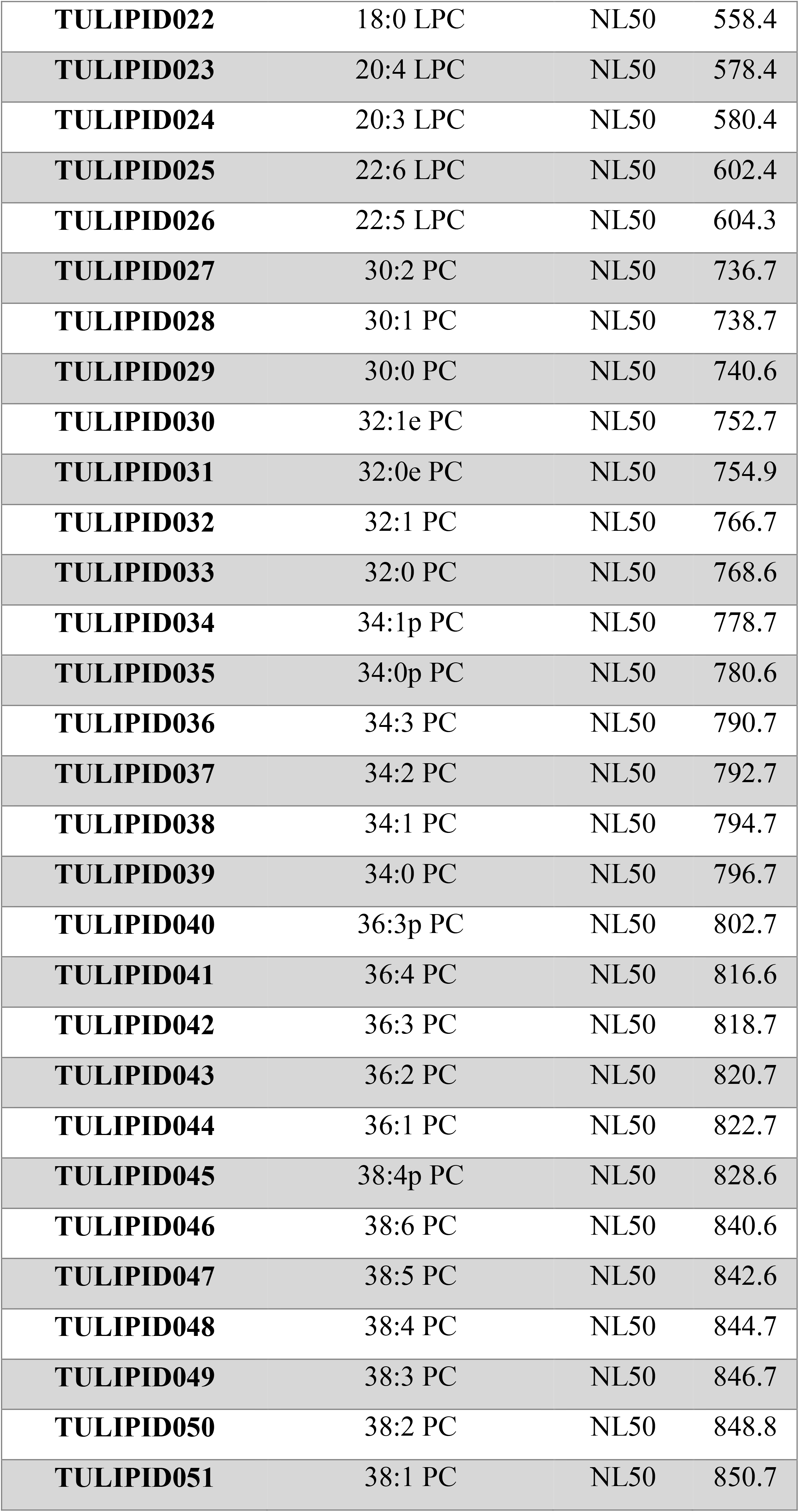
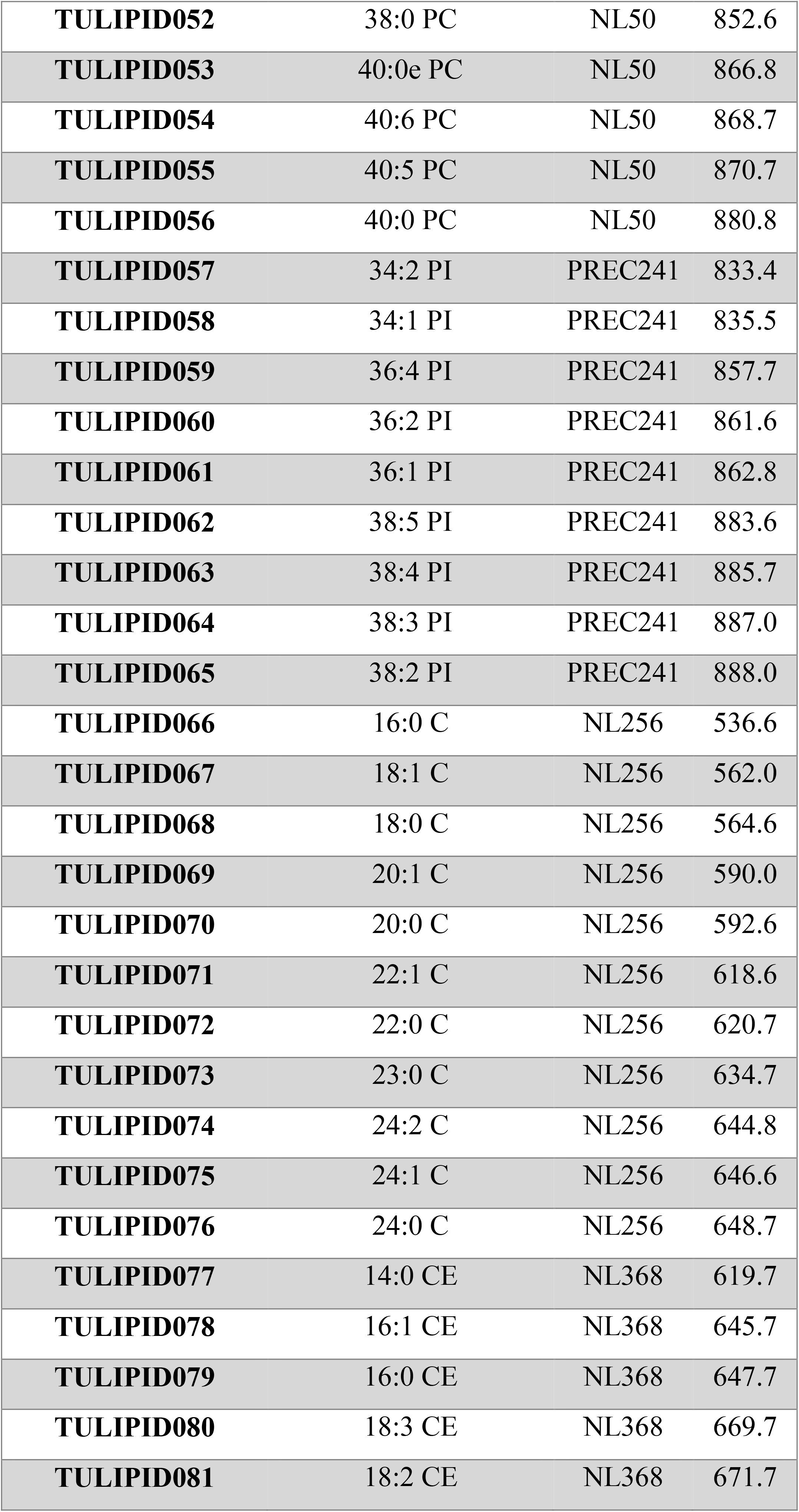
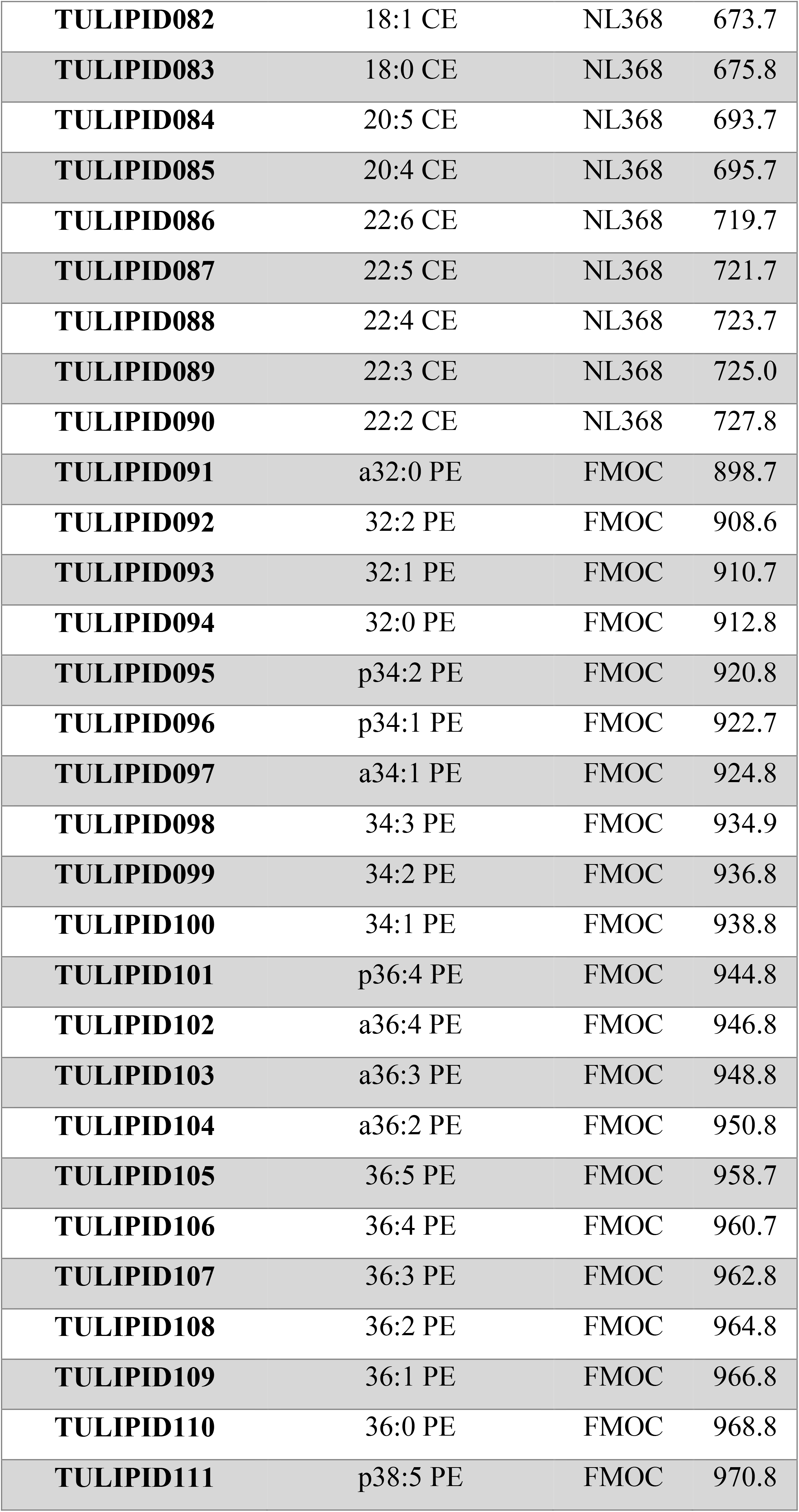
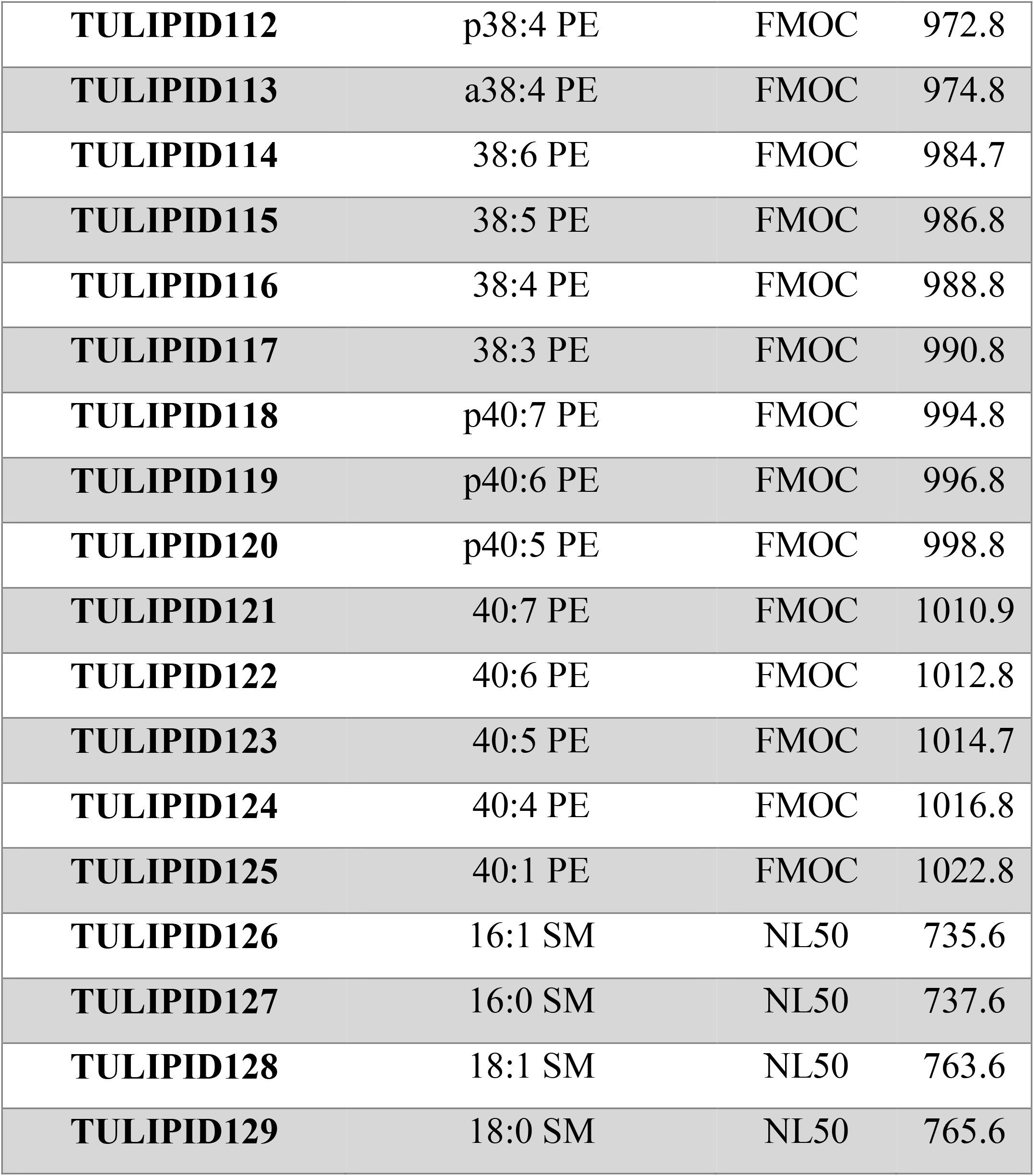
Table of positively identified targeted lipids in serum samples. A list of all of the identified targeted lipids are shown in this table with corresponding mass to charge (m/z) value and lipid scan type/experiment.

